# Development of a species diagnostic molecular tool for an invasive pest, *Mythimna loreyi* using LAMP

**DOI:** 10.1101/2020.10.01.323089

**Authors:** Hwa Yeun Nam, Min Kwon, Hyun Ju Kim, Juil Kim

## Abstract

The *Mythimna loreyi* (Duponchel) is one of the well-known a noctuid pest in Africa, Australia, and many Asian countries. This species has recently emerged as an invasive pest of some cereal crops in Korea. However, it is extremely difficult to identify the morphologically similar species, *Mythimna separate*, which occur at the cornfield in the larvae stage. Therefore, it is hard to accurately investigate invasive pests. In this study, the LAMP assay was developed for rapid, simple, effective species identification. By analyzing the mt genome, the species-specific sequence was found at the coding region of the NADH dehydrogenase subunit 5 gene. Based on this unique sequence, four LAMP primers and two loop primers were designed. The F3 and B3 primers were able to diagnose species-specific in general and multiplex PCR, and specifically reacted within the inner primers in LAMP assay. The optimal incubation condition of the LAMP assay was 61 □ for 60 minutes with four LAMP primers, though additional loop primer, BF and LF, did not significantly shorten the amplification time. The broad range of DNA concentration was workable in LAMP assay, in which the minimum detectable DNA concentration was 100 pg. Here, DNA releasing method was applied which took five minutes of incubation at 95 □ without the DNA extraction process, and only some pieces of tissue from larvae and adult samples were needed. The incidence of invasive pests is gradually diversifying, therefore, this simple and accurate LAMP assay possibly applied in the intensive field monitoring for the invasive pests and integrated management of *Mythimna loreyi*.

## 1 INTODUCTION

The *Mythimna loreyi* (Duponchel) (often called the cosmopolitan) is a noctuid pest of grain crops found in Africa, Australia, the Near East, and the Middle East and undergoes multiple generations per year (CABI, 2020). *M. loreyi* feeds on various host plants include rice, wheat, maize, sugarcane, barley, sorghum, and others, which have a large effect on female fecundity. The fecundity of female moths is greatest when the larvae feed on maize in Egypt (El-Sherif, 1972). Since *M. loreyi* is facilitated to breed, some researchers focus on the identification of products secreted by the adult corpora allata (Ho et al. 1995). Not only physiological understanding of this species (Hsieh et al., 2001; Hsieh et al., 2002; Kou, 2002; Kou and Chen, 2000) but also ecology-based developmental characteristic (Qin et al., 2017) and flight performance (Qin et al., 2018) was also studied. For the biological control, *M. loreyi* densovirus (MIDNVs) isolated in Egypt and characterized (El-Far et al., 2004; Fediere et al., 2004).

The outbreak of this pest and damage on crops has been proliferated particularly in some Asian countries. In Japan, *M. loreyi* typically occurs together with *M. separate* (Walker) that has significantly negative affects crop production (Hirai, 1975). Also, *M. loreyi* has begun to occur and damage host plants together with *M. separate* (Guo *et al.*, 2003). During a couple of years in 2019 and 2020, there have been reports that cornfield has been damaged by larvae of *M. loreyi* in Korea. This indicated that there is a possibility that *M. loreyi* can change into a sporadic pest which can cause serious damage to crops. As per these cases, *M. loreyi* occasionally damaged with its sister species, *M. separate*. However, it is difficult to distinguish two species at the cornfield in the larvae stage.

Only one molecular diagnostic tool has been studied to distinguish the *M. loreyi* which are based on the sequencing of part of the mitochondrial COI gene (Jinda, 2019). However, only one mutation existed within the 658 bp amplicon. Moreover, high sequence similarity showed between *M. loreyi* and *M. separata* which indicates the limitation of diagnosis of this pest. Therefore, we developed a simpler technique, termed loop-mediated isothermal amplification assay (LAMP), is also widely used for the rapid and accurate identification of pest species (Blaser et al., 2018; Hsieh et al., 2012; Y. H. Kim et al., 2016). The LAMP is a rapid, simple, effective, and specific amplification of DNA compared to real-time PCR based on the mitochondrial gene. It is performed under isothermal conditions that require a set of four primers, a strand-displacing DNA polymerase, and a water bath or heat block to maintain the temperature at about 65 °C following a one-time denaturation at 95 °C (Notomi et al., 2000) or one step incubation at about 65 °C (Nagamine et al., 2002).

Following the first infestation of *M. loreyi* in Korea, there is great demand from agricultural research, extension services, and farmers for diagnostic methods for these species. Therefore, we present a method based on LAMP-based on specimens collected in Korea and other sequences from GenBank. This method should be useful in assisting the effective pest management of *M. loreyi*.

## 2 MATERIALS AND METHODS

### 2.1 Sample collection and mitochondrial genome sequencing

The larval stage of *Mythimna loreyi* Korean populations was collected from Hadong (35°02.17 N, 127°47.12 E) in cornfield, 2019. Some larvae reared in the lab for morphological conformation in the adult stage and some of the individual larva directly genomic DNA was extracted with DNAzol (Molecular Research Center, Cincinnati, OH) and quantified by Nanodrop (NanoDrop technologies, Wilmington, DE, USA). Species were identified via sequencing using universal primers, LCO1490 and HCO2198 (Folmer et al., 1994). For mitochondrial genome sequencing, the Miseq platform was used and more than 1 Gb were sequenced. To assemble these data, the CLC Assembly Cell package (version 4.2.1) was used. After trimming raw data using CLC quality trim (ver. 4.21), the assembly was accomplished using the CLC de novo assembler with dnaLCW. Assembled sequences were confirmed by BLASTZ (Schwartz et al., 2003). The GeSeq program was used for annotation (Tillich et al., 2017) and the result manually checked based on the alignment of other Noctuidae species mitochondrial genomes using MEGA 7 (Kumar et al. 2016).

### 2.2 Phylogenetic analysis and primer design

Molecular phylogenetic analysis of mitochondrion genomes was inferred by using the maximum likelihood method implemented by MEGA 7 with bootstrapping (Kumar et al. 2016; Sanderson and Wojciechowski. 2000). Mitochondrial genome sequences of other Noctuidae species downloaded from GenBank, NCBI. For comparative analysis, mitochondrial genomes were aligned using mVISTA (Frazer et al. 2004; Mayor et al. 2000). Based on the global alignment result, partial sequences were re-aligned for LAMP primer design using PrimerExplorer V5.

### 2.3 LAMP and PCR

WarmStart® LAMP Kit (New England Biolabs, Ipswich, MA) used for LAMP assay. The general protocol of LAMP was flowed by the manufacture’s guideline in a 25 μl reaction mixture. For the general PCR, TOYOBO KOD□FX Taq™ (Toyobo Life Science, Osaka, Japan) used in this study. Appropriate primers with the following PCR amplification protocol: a 2□min denaturing step at 94 °C and a PCR amplification cycle consisting of denaturing at 94 °C for 20 s, annealing at 60 °C for 20 s, and extension at 68 °C for 30 s, which was repeated 35 times. The amplified DNA fragments were separated using 1.5% agarose gel electrophoresis, visualized with SYBR green (Life Technologies, Grand Island, NY). Used *M. loreyi* samples were collected in a cornfield as previously described. Pheromone traps were used for adults’ sample collection such as *M separate, Agrotis segetum, Spodoptera frugiperda, S. exigua, S. litura*, and *Helicoverpa armigera*). Traps were set in Pyeongchang (37°40.53 N, 128°43.49 E), Hongchen (37°43.35 N, 128°24.33 E) and Gangneung (37°36.56 N, 128°45.59 E) (Kim et al. 2018). DNA samples were prepared using DNAzol from trapped adults. Three biological DNA samples were used in each LAMP and PCR.

## 3 RESULTS

### 3.1 Mitochondrial genome sequencing and primer design

15,320bp of mitochondrial genome verified after trimming from about 2.2 Gb (7,312,504 reads) nucleotide sequences information obtained through Miseq. The mitochondrial genome of *Mythimna loreyi* was assembled (MT506351). The mitochondrial genome includes 13 protein-coding genes: NADH dehydrogenase components (complex □, ND), cytochrome oxidase subunits (complex □, COX), cytochrome oxidase b (CYPB) and two ATP synthases; two ribosomal RNA genes and 22 transfer RNAs.

As a result of MegaBLAST, the most homologous species was an allied species, *Mythimna separate*, which showed 93.7% similarity. The genus of Spodoptera (CABI, 2020), such as *Spodoptera exigua, S. litura, S. frugiperda*, which are possibly found together with *M. loreyi* in the cornfields showed about 89 to 90% homology based on mt genome sequence. *Agrotis segetum* (Erasmus et al., 2010), which in particular occurs and damage at the corn seedling stage, showed 90.6% similarity to *M. loreyi* based on mt genome sequence (data not shown).

The phylogenetic relationship between 15 mt-genomes of 14 species was examined (Fig. 1A), to verify a specific nucleotide sequence that only *M. loreyi* possessed among the related species with high gene similarity or a morphologically similar pest. The phylogenetic relationship result was almost similar to the megaBLAST result. The result of the phylogenetic relationship was mostly similar to the megaBLAST result. Based on the mVISTA alignment results, conserved regions among Noctuidae species and variable regions were observed (Fig. 1B). By combining the two results, the partial sequence in 8 species of 9 ND5 mt-genome was re-aligned to design the specific primer of the *M. loreyi*. Finally, four essential primers and two loop primers were designed (Fig. 2 and Table 1). Among the six primers, F3 is the specific primer that enables the diagnosis of *M. loreyi*, and only *M. loreyi* had the CCCC sequences in 4 priming regions, which are marked in the red box. A total of 4 populations that were collected from different regions had the same nucleotide sequence. Therefore, it could be sufficiently used for species-diagnosis. The basic species-diagnosis primer production strategy is the same as the previously reported species diagnosis development method of *Spodoptera frugiperda* (Kim et al., 2020).

**Fig. 1.**
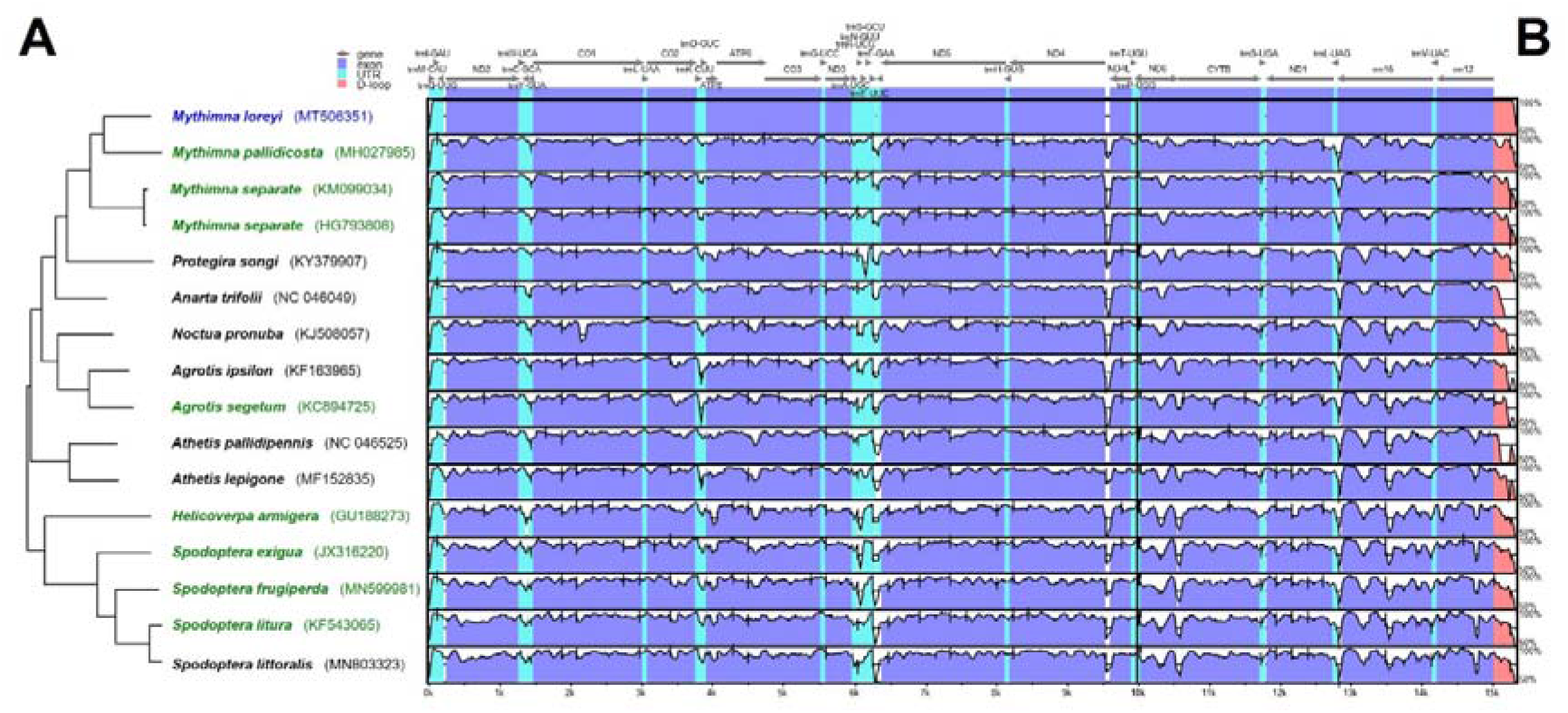
Comparison of entire mitochondrial genomes of some Noctuidae pests including newly sequenced *Mythimna loreyi*. (A) Phylogenetic relationship inferred using maximum likelihood under MEGA7. (B) Schematic diagram of the genes and their flanking regions showing the sequence diversity in mVISTA. UTR, D-loop denotes untranslated region and displacement-loop, respectively. Eight green colored mt genome sequences were re-aligned for primer design with that of target species, *M. loreyi* (Fig. 2).

**Fig. 2.**
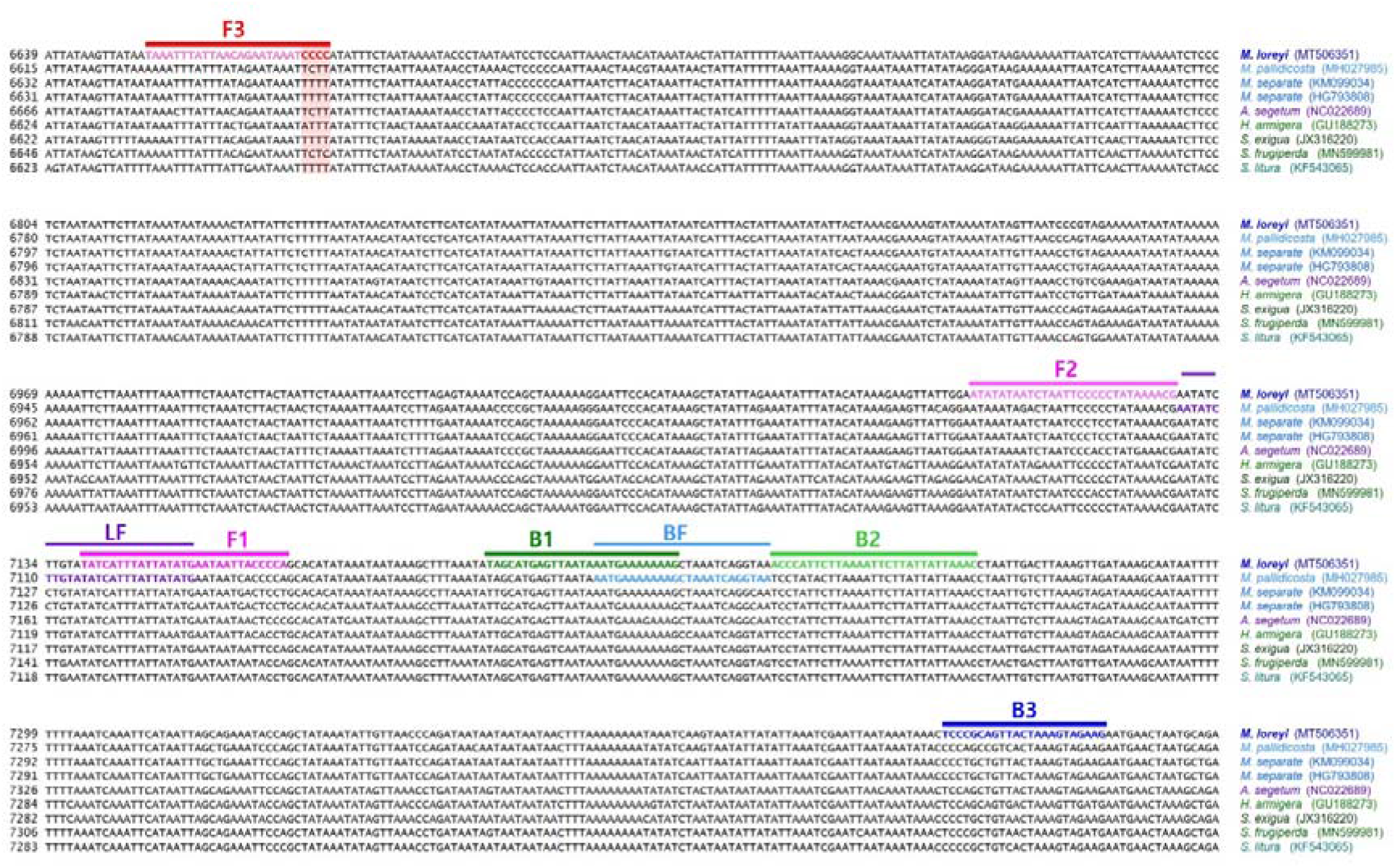
Location of primers and primer binding regions on partial sequence of some Noctuidae pests’ mtDNA for species identification of *M. loreyi*. Inner primer, FIP is consists of F1c (complementary sequences of F1) and F2. Another inner primer, BIP is also composed of B1 and B2c (complementary sequences of B2). Essential four LAMP primers (F3, FIP, BIP, and B3) generate the dumbbell structure, and two loop primers, LF and LB accelerate the LAMP reaction (see Nagamine et al. 2002 for details). Used primer information is documented in Table 2.

**Table 1.**
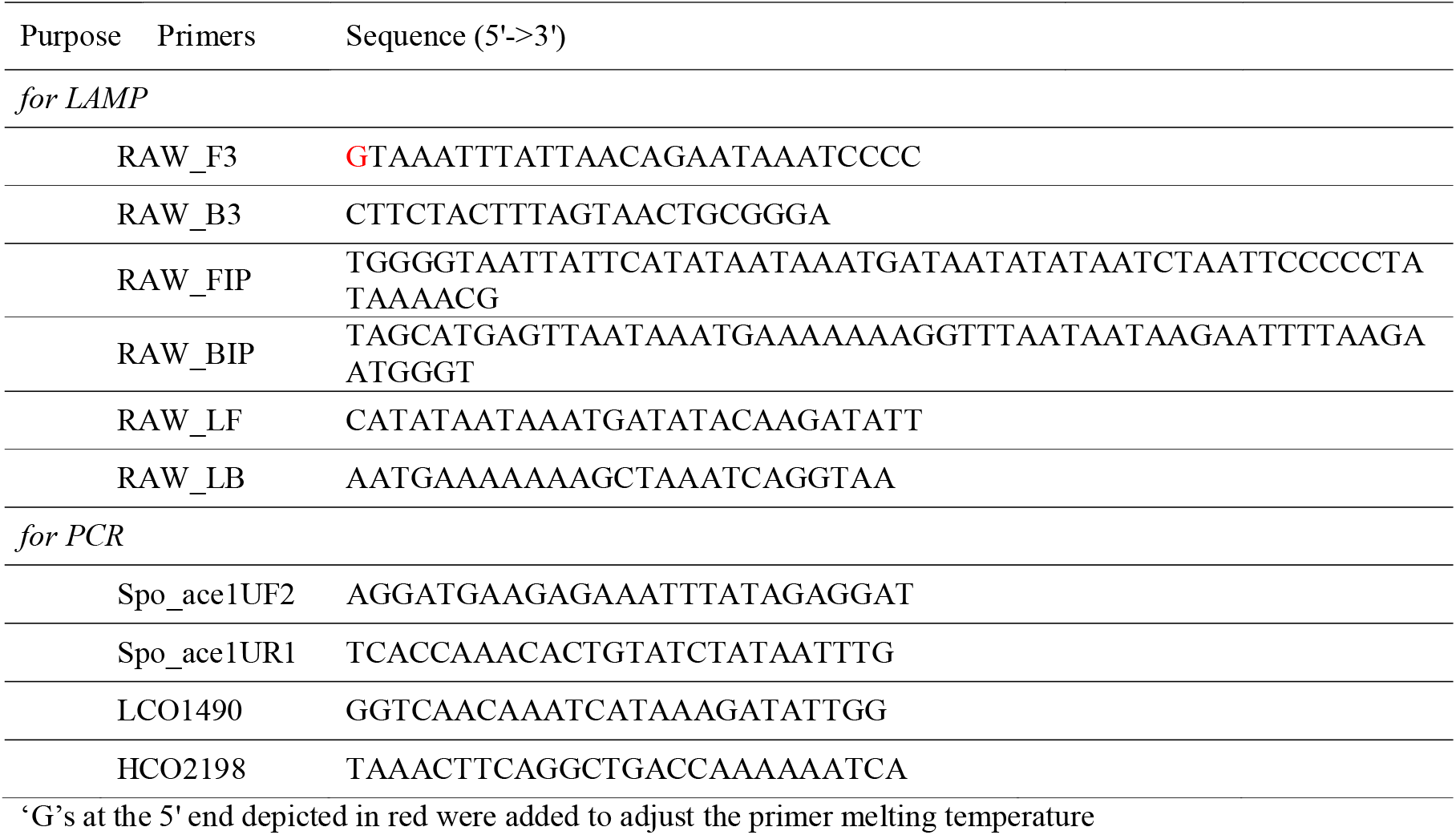
Primer list for LAMP (lamp loop mediated isothermal amplification) and PCR in this study

### 3.2 Diagnostic LAMP and PCR

As previously reported, the sensitivity of LAMP may vary depending on the temperature and reaction time (Kim et al., 2020), therefore the reaction was performed at 65, 63, and 61 □ to find the optimal reaction conditions (Fig. 3). As the amount of template DNA was quantified as 50 ng and reacted at each temperature with 25ul reaction volume, the diagnosis result confirmed in 120 min at 65 □ and 90 min at 63 □. The diagnostic level of reaction did not occur when the reaction performed less than the corresponding time in each temperature. Despite the relatively low temperature, at 61 □, the diagnostic level of reaction was confirmed in only 60 min and the false-positive reaction did not appear only once in the results of more than 3 repetition tests. Despite the relatively low temperature, at 61 □, the diagnostic level of reaction was confirmed in only 60 min, and the false-positive reaction did not appear only once in three repetition tests (Fig. 3C). Under 100 bp sized bands are not the LAMP product. That’s aggregated primers. So that generated even in the negative control.

**Fig. 3.**
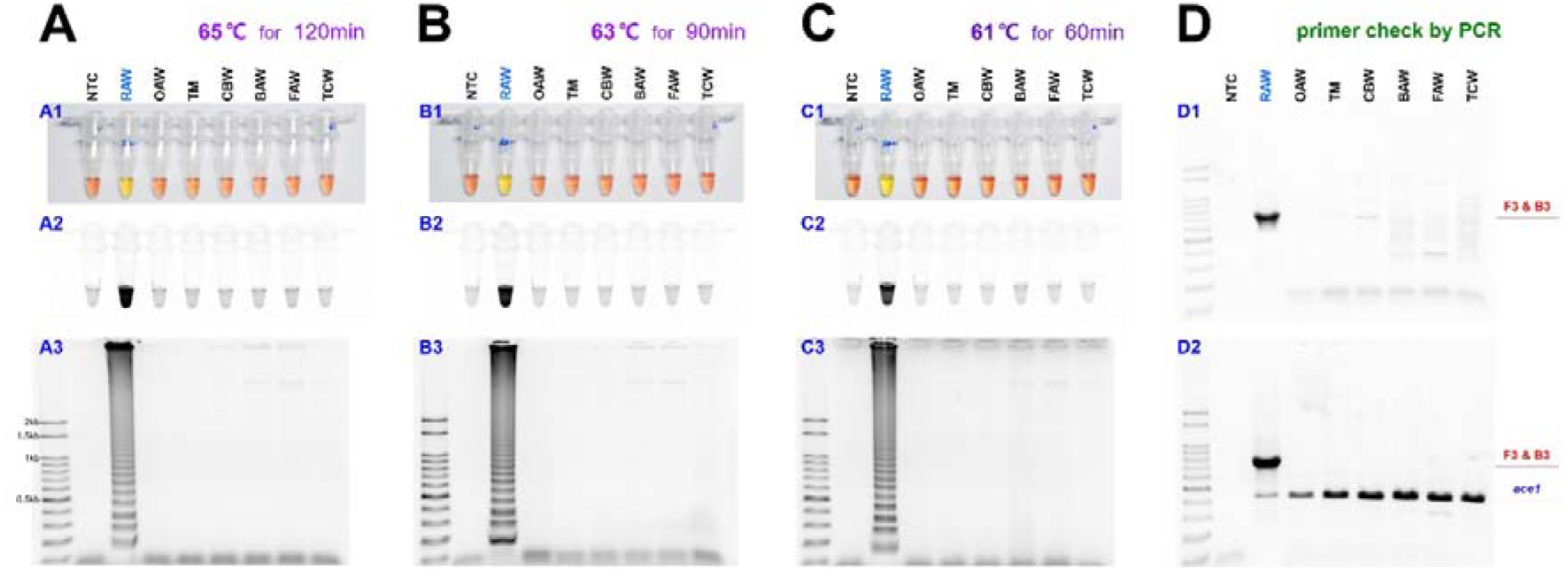
The sensitivity of the LAMP assay results in three temperature conditions such as 65, 63, and 61 °C for *M. loreyi* species detected under (A1, B1, and C1) visible light, (A2, B2, and C2) ultraviolet light with Cyber Green and (A3, B3, and C3) gel electrophoresis. The original pink color of the reaction mixture turned yellow in a positive reaction when the product was formed but remained pink in negative reactions. (D) Conventional and multiplex PCR to distinguish *M. loreyi*. 794bp amplicon amplified only in *M. loreyi* and conserved partial sequence of *ace1* type acetylcholinesterase gene was targeted as an internal reference. Abbreviations are NTC (non-template control), RAW (rice armyworm *Mythimna loreyi*), OAW (oriental armyworm *M. separata*), TM (Turnip moth *Agrotis segetum*), CBW (cotton bollworm *Helicoverpa armigera*), BAW (beet armyworm *S. exigua*), FAW (fall armyworm *S. frugiperda*), and TCW (tobacco cutworm *S. litura*)

In the LAMP assay, to confirm the diagnosis primer F3, it was reacted with reverse primer B3 through PCR, and 794bp of PCR product was identified (Fig. 3D1). Also, a species-specific reaction was confirmed, as we performed the multiplex PCR with a universal positive control primer set that targets *ace1-type* acetylcholinesterase, which can produce a positive reaction in all samples (Fig. 3D2). The two-loop primers were tested for possible enhancement of the LAMP reaction, as suggested by Nagamine et al. (2002) (Nagamine et al., 2002). As a result of reacting two-loop primers with each or two together at 61 □ for 60 min, there was no difference between the reaction result of using only four primers and shortening the reaction time (Fig. 4). Even when 10ul of the reaction solution was verified by electrophoresis after the LAMP reaction, there was no difference in band intensity. It was possible to reliably diagnose up to 100 pg when reacting using 4 LAMP primers without adding a loop primer (Fig. 5A). Therefore, *M. loreyi* species-diagnosis LAMP method that we developed can diagnose under various DNA concentration conditions from 100ng to 100pg. To increase the usability in the field, we cut a part of the tissue of the adult antenna or leg and put it in 30ul distilled water, and react at 95 □ for 5 minutes that template DNA secured without a separate DNA extraction process (Fig. 5B4). As we measured each sample of DNA concentration obtained through this DNA releasing method, each sample showed various measurements. It is possible to specific-species diagnosis in the same method of using the template DNA, which was obtained through separate DNA extraction as a positive control since it was in the range of LAMP reaction (Fig. 5B).

**Fig. 4.**
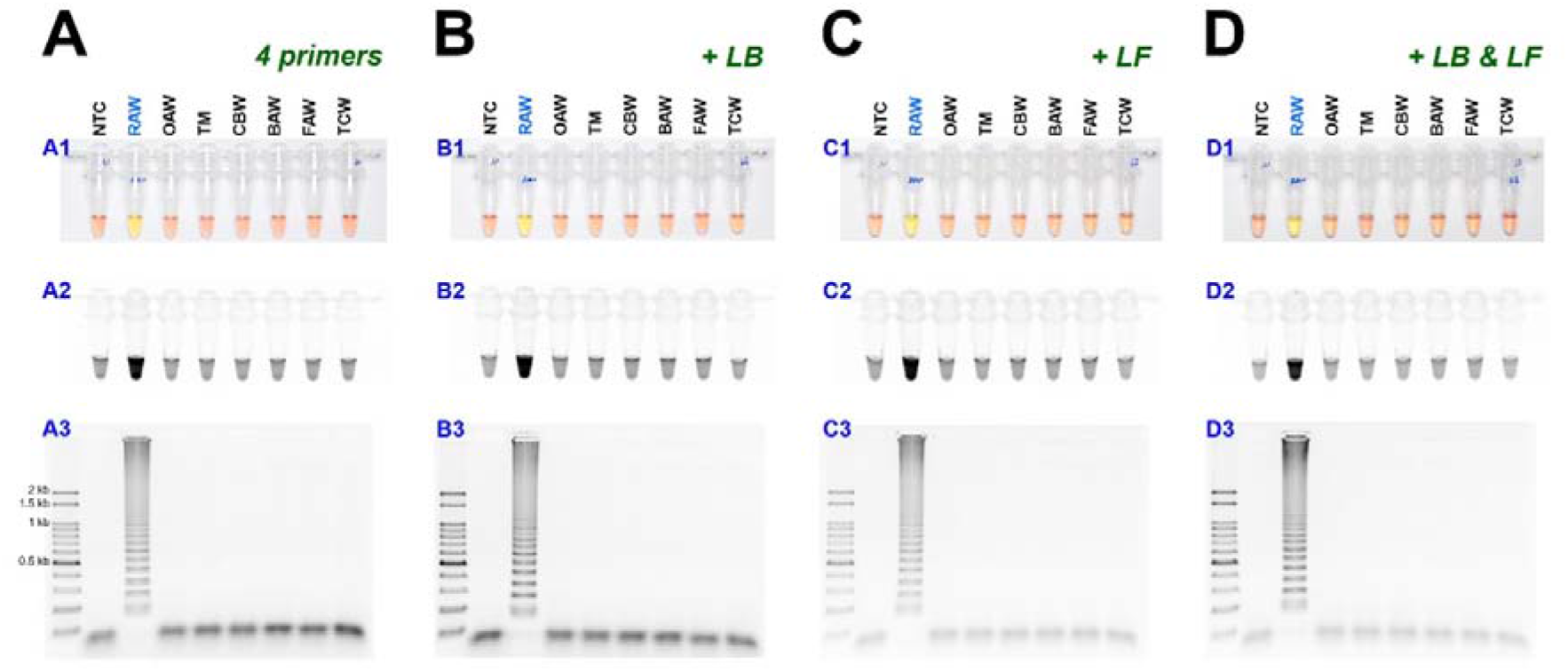
The LAMP assay results with (A) 4 primers and additional loop primers, (B) loop forward, LF, or (C) loop backward, LB, or (D) two-loop primers, LF and LB under visible light, ultraviolet light with Cyber Green and gel electrophoresis. Abbreviations as in Fig. 3.

**Fig. 5.**
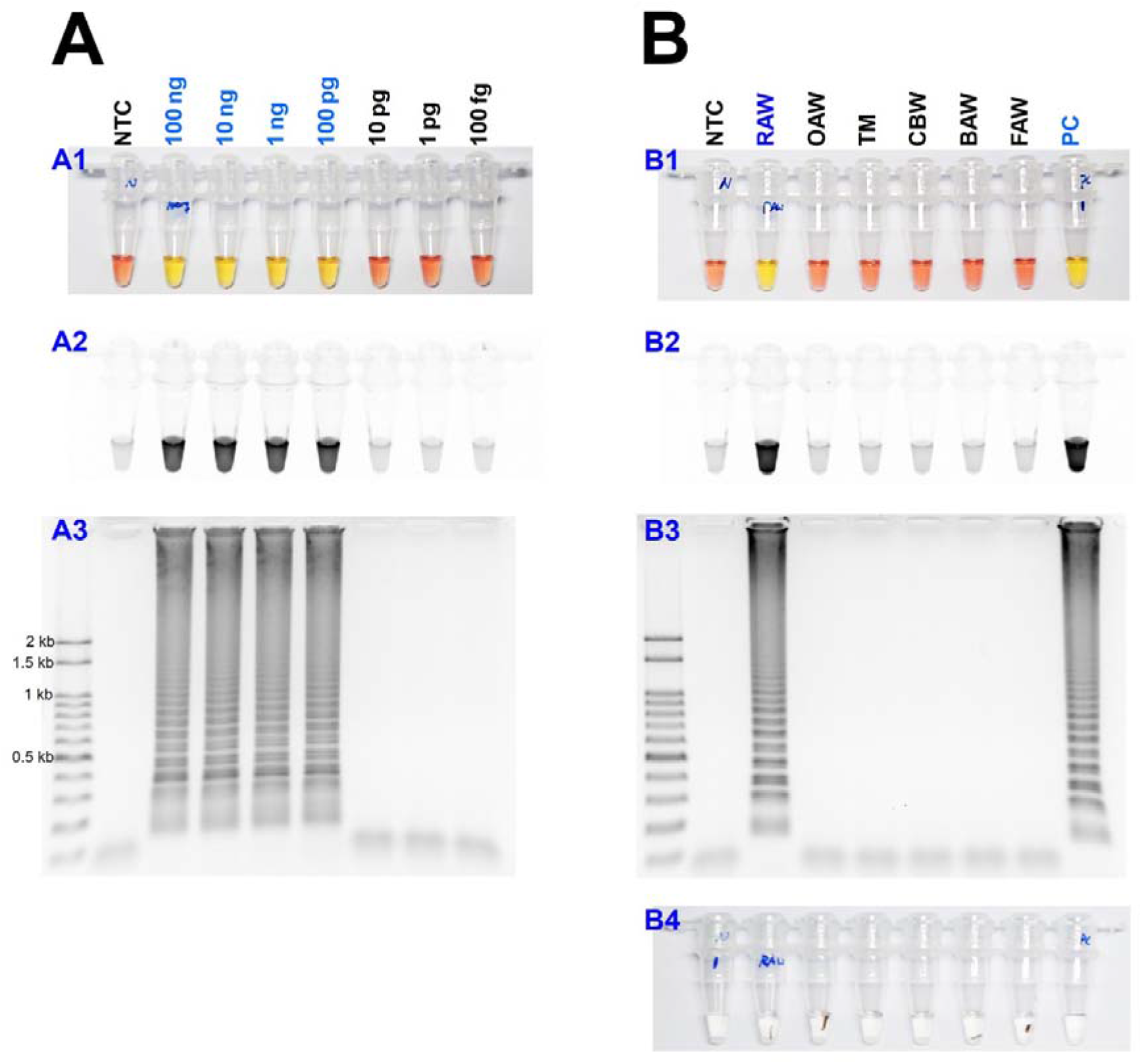
(A) Identification of the detection limit of genomic DNA in the LAMP assay from 100ng to 100fg under visible light, ultraviolet light with Cyber Green, and gel electrophoresis. (B) The sensitivity of the LAMP assay results with the DNA releasing technique from insect tissue. Around 10mg of the adult leg (or antenna) were incubated at 95 □ for 5minutes (B4). Abbreviations as in Fig. 4, except PC (positive control, isolated DNA from *M. loreyi*).

## 4 DISCUSSION

Invasive pests such as *S. frugiperda* are increasing worldwide due to global warming and climate change (Li et al., 2020; Goergen et al., 2016). Also, *M. loreyi* is a species that originates from China and damage from it gradually spreading. As with other invasive pests found in the field, they are often similar in morphology to allied species, which are difficult for an investigation of initial density and pest management. Species diagnosis methods for invasive pests such as *S. frugiperda* have been developed and utilized for this reason (Kim et al., 2020). Currently, the molecular biology method for diagnosing *M. loreyi* species is only using the mtCO1 universal primer (LCO1490, HCO2198), and process PCR and sequencing to compare the degree of homology. However, this method requires a lot of time and effort, such as DNA extraction, PCR, electrophoresis, and sequencing. Therefore, we developed a molecular diagnosis method that can diagnose species within a short time without a separate DNA extraction process, and only a heat block is needed that can control temperature (Fig. 5). The basis diagnosis strategy is very similar to the *S. frugiperda* species diagnosis method (Kim et al., 2020). This method can complete all experimental procedures and verify the results within 1 hour and 30 minutes right after obtaining a sample. In this study, only the results specified adult samples, but it is possible to use larvae.

The simplicity, accuracy, and adaptability for high throughput of the LAMP assay are distinct advantages (Mori and Notomi, 2009; Notomi et al. 2015; Zhang et al. 2014). Moreover, recently LAMP utilized in various fields, such as many ecology studies, medical aspects, outside of the lab, and can be applied to diagnose plant viruses in insect body and insecticide-resistant gene mutation (Lee, 2017; Choi et al., 2018). Also, the diagnostic primer used for LAMP can be used for various diagnostic methods, because it was possible to apply in general PCR and multiplex PCR (Fig. 3). Therefore, it is feasible to diagnose a larger sample with positive control in the form of multiplex PCR, which sufficiently modified and used in the laboratory. There are advantages and disadvantages to find a species diagnostic marker in the mt genome as well as in part of the genomic DNA. First of all, the disadvantage is that LAMP primer production is limited because there are many parts of AT-rich. In advantage, the number of copies is large that possible to diagnose with a small amount of DNA or DNA releasing method. On the contrary, a species diagnosis marker in genomic DNA should be found in exon rather than intron because it has a large variation. But it is difficult to develop marker in exon because often quite conserved within an allied species. Also, the DNA releasing method can be used, but the efficiency is low when the copies of the gene are small (personal communication). Therefore, in this study, a species diagnosis marker was designed in the mt genome to combine with a DNA releasing method that is highly applicable in the field. Therefore, in this study, a species diagnosis marker was designed within the mt genome to combine it with a DNA releasing method that is highly applicable in the field. Moreover, a significantly efficient method was developed, targets the *ace1* gene as a positive control of the species to be compared. This simple and accurate diagnosis using LAMP assay possibly applied in the intensive field to monitor and pest management of *M. loreyi*.

## ACKNOWLEDGEMENTS

The Cooperative Research Program supported this study for Agriculture Science & Technology Development (Project No. PJ01509301), the Rural Development Administration, Republic of Korea.

## AUTHORS’ CONTRIBUTIONS

HYN and JK conceived the study. HYN, MK, HJK and JK prepared samples and performed mt genome sequencing. HYN and JK performed experiments and mainly wrote the paper. All authors read and approved the manuscript.

